# Association analysis of repetitive elements and R-loop formation across species

**DOI:** 10.1101/2020.11.09.374124

**Authors:** Chao Zeng, Masahiro Onoguchi, Michiaki Hamada

**Author notes:** Correspondence may be addressed to CZ or MH.

## Abstract

Genomes are known to have a large number of repetitive elements. Emerging evidence suggests that these non-coding elements may play an important regulatory role. However, few studies have investigated the effect of repetitive elements on R-loop formation. In this study, we found different repetitive elements related to R-loop formation in various species. By controlling length and genomic distributions, we observed that satellites, long interspersed nuclear elements (LINEs), and DNAs were each specifically enriched for R-loops in humans, fruit flies, and *Arabidopsis thaliana*, respectively. R-loops also tended to arise in regions of low-complexity or simple repeats across species. We also found that the repetitive elements associated with R-loop formation differ according to developmental stage. For instance, LINEs and long terminal repeats (LTRs) are more likely to contain R-loops in embryos (fruit fly) and then turn out to be low-complexity and simple repeats in post-developmental S2 cells. Our results indicate that repetitive elements may have species-specific or development-specific regulatory effects on R-loop formation. This work advances our understanding of repetitive elements and R-loop biology.

## Introduction

An R-loop is a three-stranded structure composed of a DNA:RNA hybrid with a displaced single-stranded DNA (ssDNA). Although R-loops were initially considered to be rare by-products of transcription, recent reports have suggested that R-loops are widely distributed in eukaryotic genomes^1–3^ and are involved in gene regulation and genome integrity^4–9^. R-loops regulate gene expression through a variety of molecular mechanisms. For instance, in promoter regions, R-loops promote or inhibit gene transcription by decreasing methylation^10^ or enhancing polycomb-mediated gene silencing^11^. Surprisingly, a recent report indicated that R-loops can shield the ribosomal gene expression by RNA polymerases II from the transcription conflicts caused by other RNA polymerases^12^. In terminator regions, R-loops can induce RNA polymerase stalling to improve the efficiency of transcription termination^13^. Simultaneously, these R-loops will promote nascent RNA cleavage and 3 transcript degradation^14^. With respect to genome integrity, R-loops can affect genome dynamics^15^ and telomere stability^16^. In centromeric regions, R-loops are reported to maintain genome stability by promoting chromatin condensation^15^ or chromosome segregation^17^. In telomeric regions, telomere repeat-containing RNAs (TERRAs) preferentially accumulate in short telomeres to form telomere-repairing R-loops^16^. Previous reviews described other details of R-loops and genome integrity^4–9^. Additionally, with an increasing number of R-loop-related diseases (such as neurological diseases and cancers) being reported and studied^18–22^, a deep understanding of the biology of R-loop structures is critical.

Genome-wide mapping of R-loop structures, performed with immunoprecipitation-based high-throughput sequencing^3,23^, provides a global approach for quantitative analysis and systematic characterization of R-loops. This technique involves immunoprecipitating R-loop structures using a DNA-RNA hybrid antibody (S9.6) and then subjecting those R-loop-forming DNAs to direct DNA-sequencing, known as DRIP-seq (DNA–RNA immunoprecipitation followed by high-throughput DNA sequencing)^23^. Additionally, other R-loop profiling methods have been developed in the past decade by modifying antibodies and/or protocols^2,24–29^.

Previous studies have uncovered several characteristics of R-loop formation. For example, within the genome, R-loops are prone to be formed in GC-skew (asymmetric strand distribution of guanines and cytosines)^23,30^ or AT-skew (asymmetric strand distribution of adenines and thymines)^1,2^ regions. The stability of G quadruplex (G4) structures is also related to R-loop formation^31^. Notably, multiple reports have indicated that R-loop formation is associated with short tandem repeats, especially trinucleotide repeat (TNR) expansion^32–35^. An R-loop predictive model was designed with the feature of short tandem repeats in mind^36^. Additionally, limited studies have shown that transposable elements (TEs), including TY1^37^ and LINEs^38^, may play a role in the formation of R-loops. Specific epigenetic signatures also connect with R-loop structures^3,38^. Although R-loops are thought to form co-transcriptionally or in cis (transcription and R-loop formation at the same locus)^6^, accumulating evidence has indicated that R-loop structures can also form post-transcriptionally or in trans (RNA transcribed from a locus hybridizes to a distal locus)^39–43^. However, our understanding of R-loop formation and their sequence characteristics in the genome is still limited.

In this study, we focused on the effects of repetitive elements on R-loop formation. By comparing the frequency of repetitive elements in R-loops and the controls, we derive the repeat class or family that is associated with R-loop formation. For this purpose, we separately prepared three groups of controls, corresponding to uniform distribution in the genome, specific length and genomic distributions, and co-transcriptional formation. The three controls may not be appropriate for all the data studied; for example, we found that more than 60% of R-loops in fruit fly do not overlap with the transcribed region (implying that the third control is not preferred). We observed different repetitive elements related to R-loop formation in various species. Based on the second control mentioned above, we discovered that satellites, LINEs, and DNAs were each enriched for R-loops in humans, fruit flies, and *Arabidopsis thaliana*, respectively. Additionally, R-loops were mainly found in regions of low-complexity or simple repeats across these three species. Interestingly, we also found that the repetitive elements associated with R-loop formation differ according to the organism’s developmental stage. For fruit flies, LINEs and LTRs are more likely to contain R-loops in embryos, which changes to R-loops being more prevalent in areas of low-complexity and simple repeats in S2 cells. To our knowledge, this is the first comprehensive analysis of the association between repetitive elements and R-loop formation. This work improves our insights into the potential functions of repetitive elements and our understanding of the biological mechanisms underlying R-loop formation.

## Results

### Different controls provide multiple perspectives on the association between repetitive elements and R-loop formation

To investigate the association between repetitive elements and R-loop formation, we asked whether there was an overrepresented or underrepresented repetitive element in the R-loops. For this purpose, we considered three controls. First, R-loops are randomly and uniformly distributed in the genome. Second, R-loops of specific lengths are distributed in specific regions of the genome. Finally, most R-loops are co-transcriptional and are more likely to form where nascent transcripts are produced^6^. Thus, we prepared a genome control, sampling control, and GRO control to calculate the enrichment of repetitive elements in R-loops under the corresponding conditions (see “Materials and Methods” for details). We observed from the DRIP-seq data that the median R-loop lengths in animals (human and fruit fly) ranged from 414 to 618 nt, whereas those in plants (*A. thaliana*) were longer, with a median of 998 nt (Figure 1A). In addition, only 5.5%–15.8% of R-loops were detected in the intergenic region (Figure 1B), which implies that the majority of R-loops preferentially form surrounding gene regions, which is consistent with previous reports^3,24,44^. In addition, R-loops formed most frequently in promoter regions across species (Figure 1B). For *A. thaliana*, in particular, up to 60% of R-loops were distributed in promoter regions, which reduces the occurrence of R-loops in other regions of the genome, especially in introns, where only 0.2% of R-loops were formed. To this end, a sampling control was randomly generated in different datasets, simulating the distribution of R-loop lengths and genomic locations.

**Figure 1.**
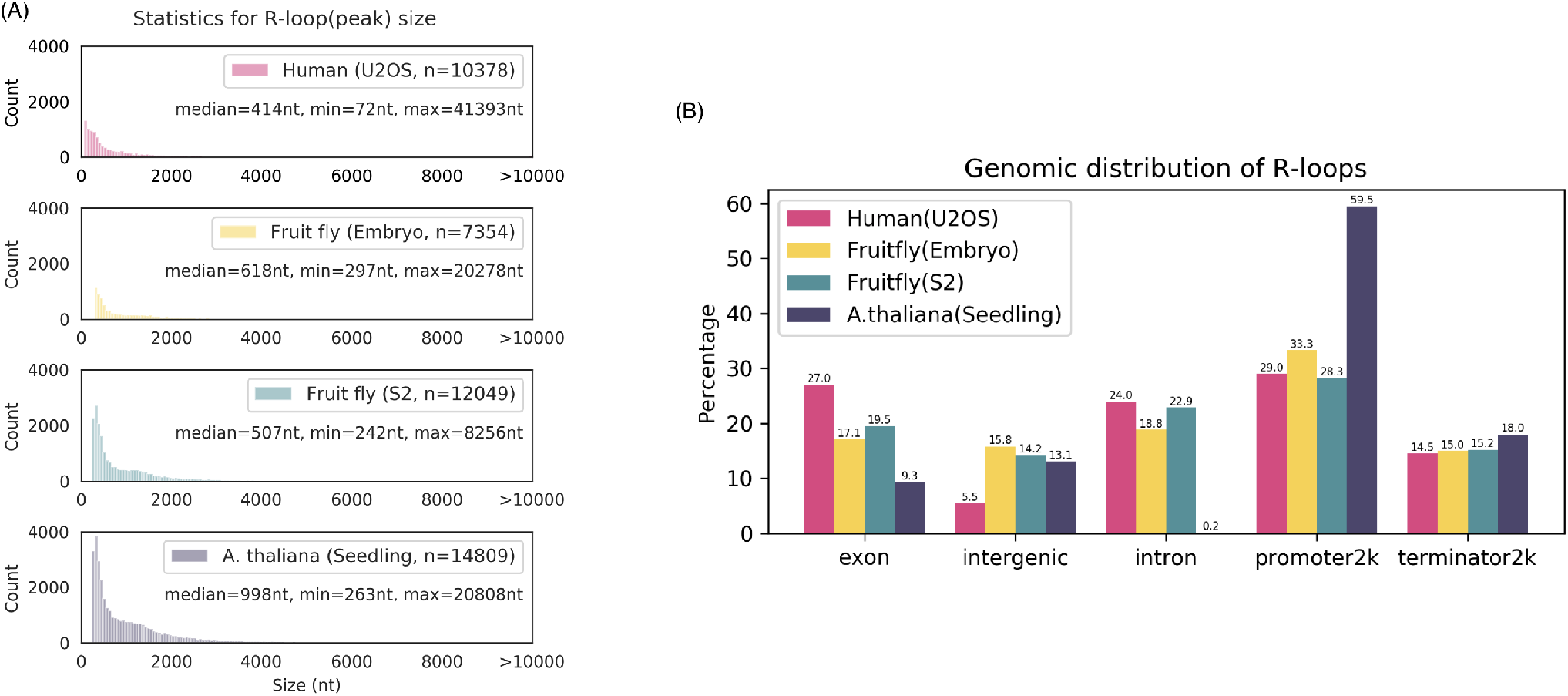
Distributions of R-loop characteristics in different datasets. (A) Size and (B) genomic distribution of R-loops. An R-loop is considered to be from a given genomic region when it overlaps more than one base with that region. In the case where an R-loop overlaps with more than one genomic region, we assigned it according to the priority (promoter2k > terminator2k > exon > intron > intergenic). Promoter2k: within 2000 nt upstream of a gene; terminator2k: within 2000 nt downstream of a gene; intergenic: regions excluding genes, promoter2ks, and terminator2ks.

Considering that R-loops may form in irregular transcribed regions (e.g., enhancers, promoters) that are not detectable by ordinary RNA-seq, we used GRO-seq data that can identify nascent transcripts to define transcriptional regions (including genes) in the genome for different tissues or cell lines. We then randomly generated the GRO control from real-time transcriptional regions. Notably, according to the GRO-seq data, the proportion of nascent transcripts detected in the intergenic region varied widely among species (3.18%–10.43%, Table S1), and was highest in humans. We speculate that the reason for this is the presence of a fraction of unannotated genes, and U2OS is a cell line that may have cancer-specific transcripts. To verify whether the majority of R-loops are co-transcriptional, we checked the percentage of overlaps between R-loops (as defined by DRIP-seq) and transcribed regions (as defined by GRO-seq). Although humans have 10.43% of transcribed regions in intergenic areas (see Table S1), the overlap between R-loops and these transcribed regions is the highest (70%) compared to the other two species (Figure 2). The lowest overlap between R-loops and transcribed regions was observed in fruit flies (embryos, 24%). It can be observed that all the transcribed regions defined in the GRO-seq data had a high match with the annotated gene regions (84.85%–91.8%, Table S1). Therefore, we speculate that the R-loops in fruit flies (embryos) may be formed by transcripts derived from distant or other chromosomal regions.

**Figure 2.**
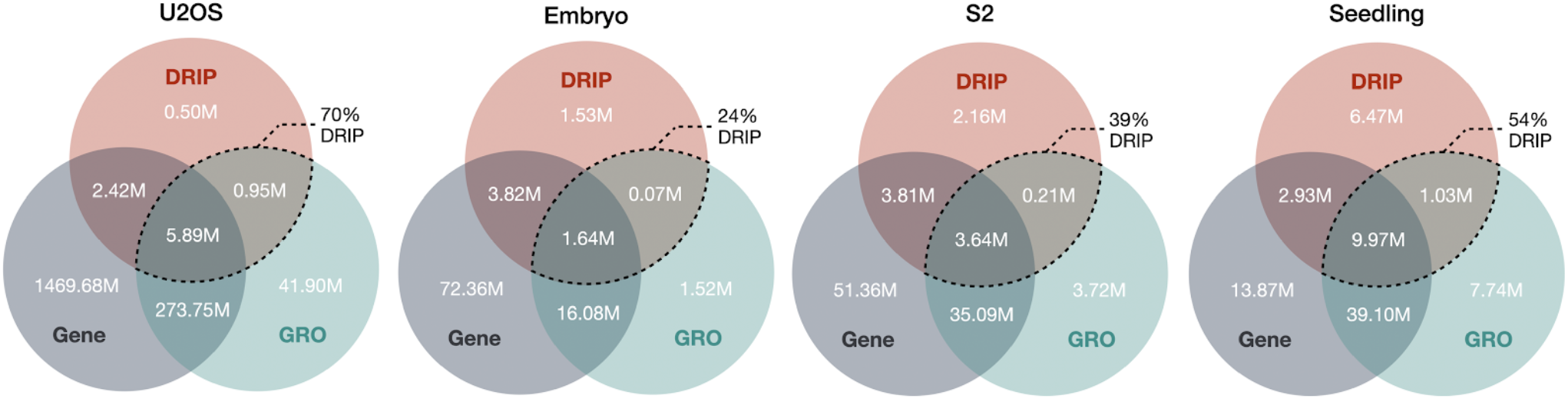
Venn diagram analysis for R-loops, transcribed regions, and genes at base level. R-loops and transcribed regions were defined by DRIP-seq (red) and GRO-seq (green), respectively. Gene regions (gray) were extracted based on gene annotations (see “Materials and Methods” for details). Areas marked with dashed lines indicate the overlap of R-loops and transcribed regions. The percentages represent the percentage of R-loops that are derived from the transcribed regions. M: million bases.

### Repetitive elements are associated with R-loop formation across species and developmental stages

By first comparing the enrichment of repetitive elements in humans, fruit flies (S2 cells), and *A. thaliana*, we observed some patterns of relative consistency among species (Figure 3, middle panel). In the sampling control, we observed that low-complexity and simple repeats tended to be enriched in R-loops among species (Z-scores were greater than 10), while RC appeared less frequently in R-loops than in random cases (Z-scores were less than −1.7). Consistent with the previous report^45^, DNA, LINEs, and LTRs were underrepresented in animal (human and fruit fly) R-loops (Z-scores between −2.25 and −23.28); however, they were overrepresented in plant (*A. thaliana*) R-loops (Z-scores between 0.56 and 24.9). In human and *A. thaliana* R-loops, short interspersed nuclear elements (SINEs) were underrepresented (Z-score was −18.78 and −2.73, respectively), whereas rRNA was mildly enriched (Z-score was 1.77 and 1.97, respectively). We also found some species-specific patterns. For example, satellites were significantly enriched in human R-loops (Z-score = 17.51), but significantly under-enriched in fruit fly R-loops (Z-score = −13.48). Notably, when we switched the controls to the transcriptional region (GRO control), we found that retroposons and satellites were enriched in human and *A. thaliana* R-loops, respectively, suggesting a positive correlation between the formation of cis R-loops and these two repetitive elements. Additionally, in humans and *A. thaliana*, the high degree of overlap between R-loops and transcriptional regions (Figure 2) enhances the confidence of the cis formation of R-loops in these two species.

**Figure 3.**
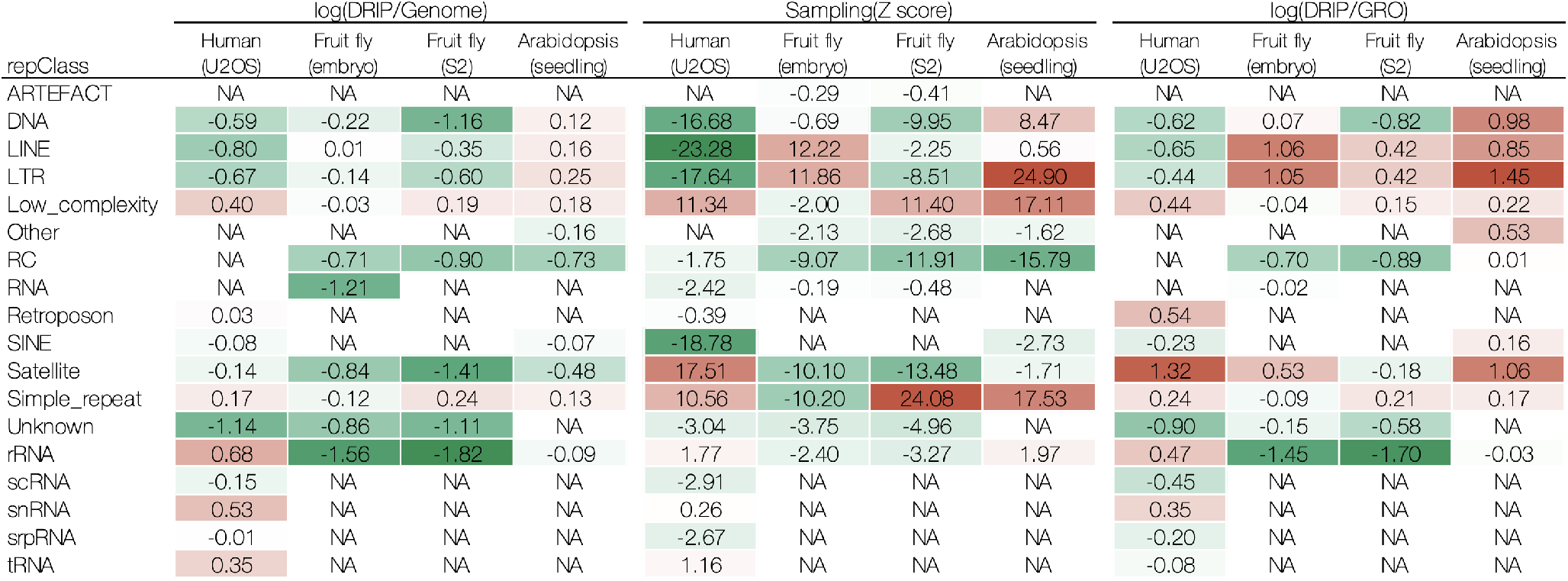
Enrichment matrix of repeat classes across human, fruit fly and *A. thaliana*. Left, middle, and right panels correspond to genome, sampling, and GRO controls, respectively. Positive: overrepresented; negative: underrepresented; NA: not available. See Figure S1 for details.

We asked whether the association between repetitive elements and R-loop formation varies during development. For this purpose, we compared embryos and S2 cells of the fruit fly. Surprisingly, in the sampling control, we found that R-loops were more likely to form in regions of the embryo containing LINEs (Z-score = 12.22) and LTRs (Z-score = 11.86) (Figure 3, middle panel). In post-developmental S2 cells, R-loops were highly aggregated in regions with low complexity (Z-score = 11.4) or simple repeats (Z-score = 24.08). However, when we applied the GRO control (Figure 3, right panel), the above results were relatively attenuated (i.e., low-complexity and simple repeats), as even LINEs and LTRs were not separated in the embryo (log(DRIP/GRO) = 1.06 and 1.05, respectively) and S2 cells (both log(DRIP/GRO) = 0.42) showing enrichment in the R-loops. Note that, in fruit flies, a small fraction (up to 39%) of R-loops overlapping with the transcribed regions might be a source of this inconsistency (Figure 2).

### Various repeat families associated with R-loop formation

Further, we investigated the relationship between R-loop formation and the repetitive elements in each species at the repeat family level. For humans, we consistently observed an enrichment of R-loops, in the sampling and the GRO controls, containing centr (Z-score = 18.96, log(DRIP/GRO) = 1.75), telo (Z-score = 20.87, log(DRIP/GRO) = 1.53), low-complexity (Z-score = 11.34, log(DRIP/GRO) = 0.44), simple repeat (Z-score = 10.56, log(DRIP/GRO) = 0.24), snRNA (Z-score = 0.26, log(DRIP/ GRO) = 0.35) and rRNA (Z-score = 1.77, log(DRIP/GRO) = 0.47) in the genome (Figure 4). Interestingly, for the GRO control, we found that SVA repeat elements (belonging to the retroposon class, log(DRIP/GRO) = 0.54) and satellites (log(DRIP/GRO) = 0.23) preferentially occurred in the R-loops.

**Figure 4.**
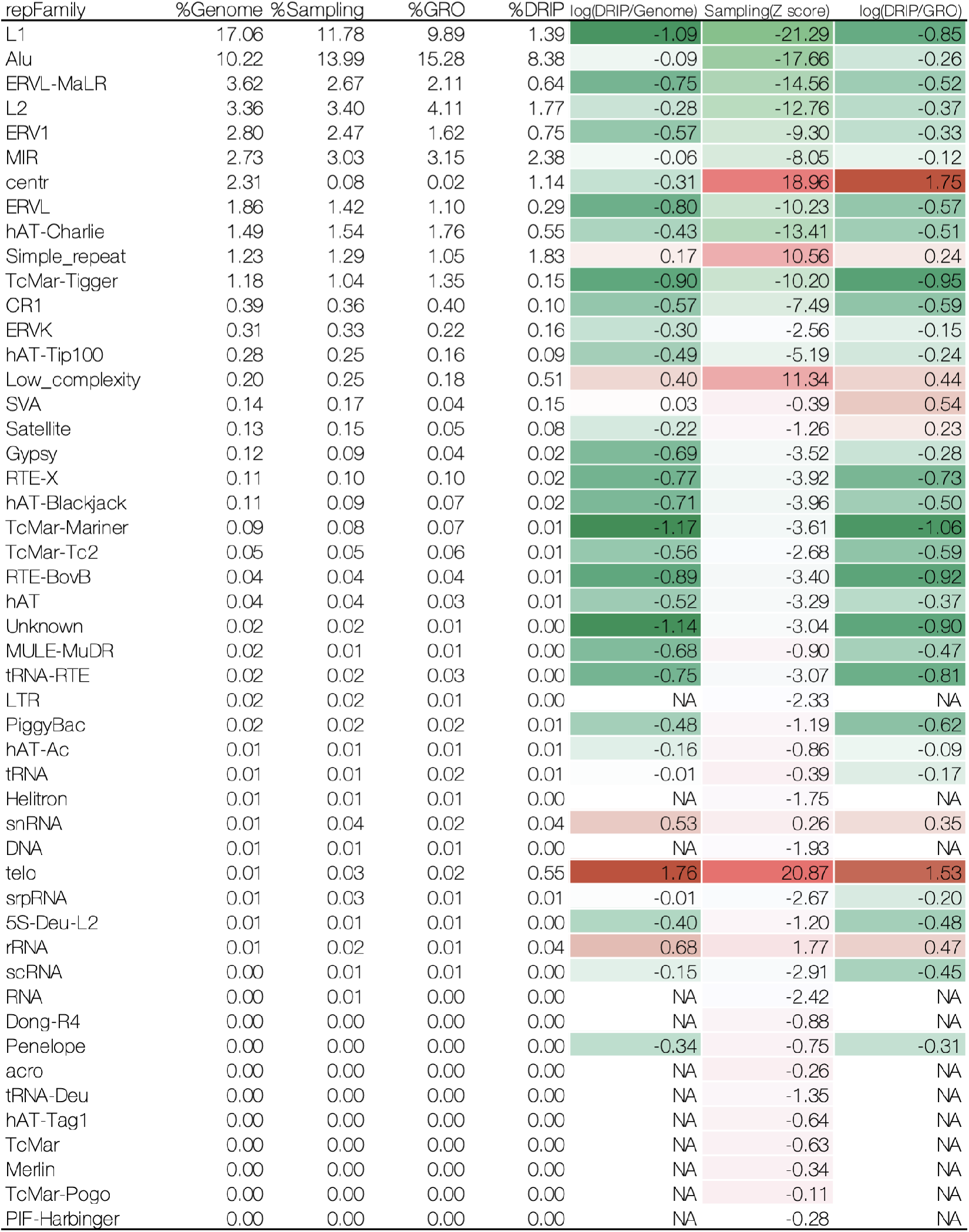
Enrichment analysis of repeat families in human U2OS cells.

For fruit fly embryos, we consistently observed the enrichment of specific repeat families in R-loops in both the sampling control and the GRO control (Figure 5). For example, Gypsy (Z-score = 3.59, log(DRIP/GRO) = 0.85) and Pao (Z-score = 16.65, log(DRIP/GRO) = 1.53), which are both LTRs; Jokey (Z-score = 5.6, log(DRIP/GRO) = 1.03), R1 (Z-score = 7.28, log(DRIP/GRO) = 1.47), CR1 (Z-score = 14.35, log(DRIP/GRO) = 0.96), and LOA (Z-score = 5.97, log(DRIP/GRO) = 0.97), all belonging to the LINEs; TcMar-Tc1 (Z-score = 6.15, log(DRIP/GRO) = 0.39) and PiggyBac (Z-score = 1.95, log(DRIP/GRO) = 0.44), both of which are in the GRO control, I (log(DRIP/GRO) = 0.23) and R2 (log(DRIP/GRO) = 1.16) belonging to L1, copia (log(DRIP/GRO) = 0.92, belonging to the LTRs, and satellites (log(DRIP/GRO) = 0.53) also tended to be enriched in R-loops. For S2 cells, the number of enriched repeat families in R-loops was significantly reduced compared to that in embryos (Figure 5). In addition to the enrichment of Pao (Z-score=2.26, log(DRIP/GRO)=0.74) and CR1 (Z-score=3.45, log(DRIP/GRO)=0.32) that can still be observed in R-loops (which is consistent with the results in embryos), we also observed an increase in simple repeats (Z-score = 24.08, log(DRIP/GRO) = 0.21) and low-complexity (Z-score = 11.4, log(DRIP/GRO) = 0.15) in the R-loops of S2 cells.

**Figure 5.**
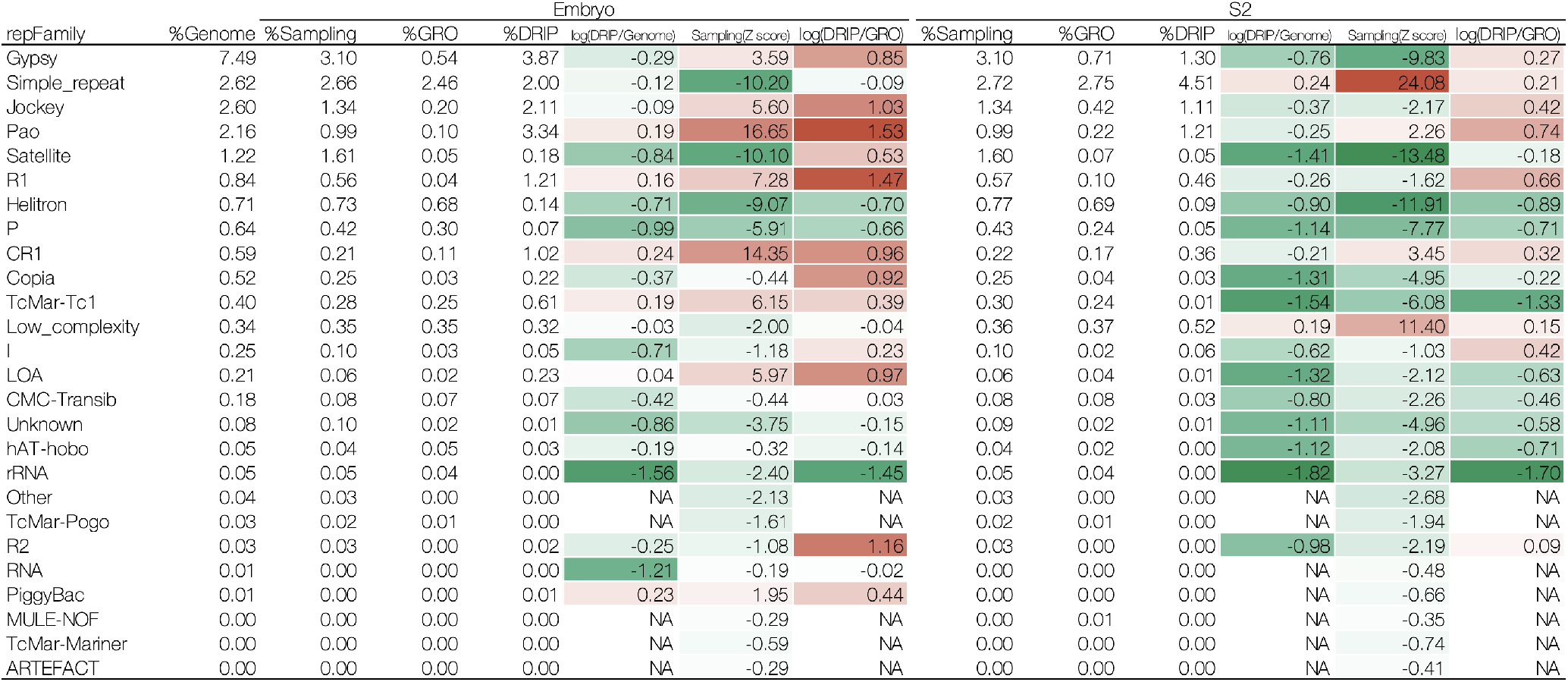
Enrichment analysis of repeat families in fruit fly (embryos and S2 cells).

For *A. thaliana*, nearly half of the repeat families were enriched in R-loops in both the sampling and GRO controls (Figure 6). These repeat families were Gypsy (Z-score = 25.7, log(DRIP/GRO) = 1.72), Copia (Z-score = 3.52, log(DRIP/GRO) = 0.99), LTR (Z-score = 8.55, log(DRIP/GRO) = 1.41), all belonging to the LTRs; MULE-MuDR (Z-score = 7.23, log(DRIP/GRO) = 1.14), CMC-EnSpm (Z-score = 7.59, log(DRIP/GRO) = 2.08), En-Spm (Z-score = 1.66, log(DRIP/GRO) = 1.52), DNA (Z-score = 0.38, log(DRIP/GRO) = 0.46), which are DNAs; L1 (Z-score = 0.56, log(DRIP/GRO) = 0.85) belonging to the LINEs; centr (Z-score = 1.34, log(DRIP/GRO) = 2.61); simple repeat (Z-score = 17.53, log(DRIP/GRO) = 0.17) and low-complexity (Z-score = 17.11, log(DRIP/GRO) = 0.22). However, PIF-Harbinger (log(DRIP/GRO) = 0.28), hAT-Ac (log(DRIP/GRO) = 0.32), all belonging to DNA, satellite (log(DRIP/GRO) = 0.58), composite (log(DRIP/GRO) = 0.53), and tRNA (log(DRIP/GRO) = 0.35), were only observed to be enriched in R-loops in the GRO control.

**Figure 6.**
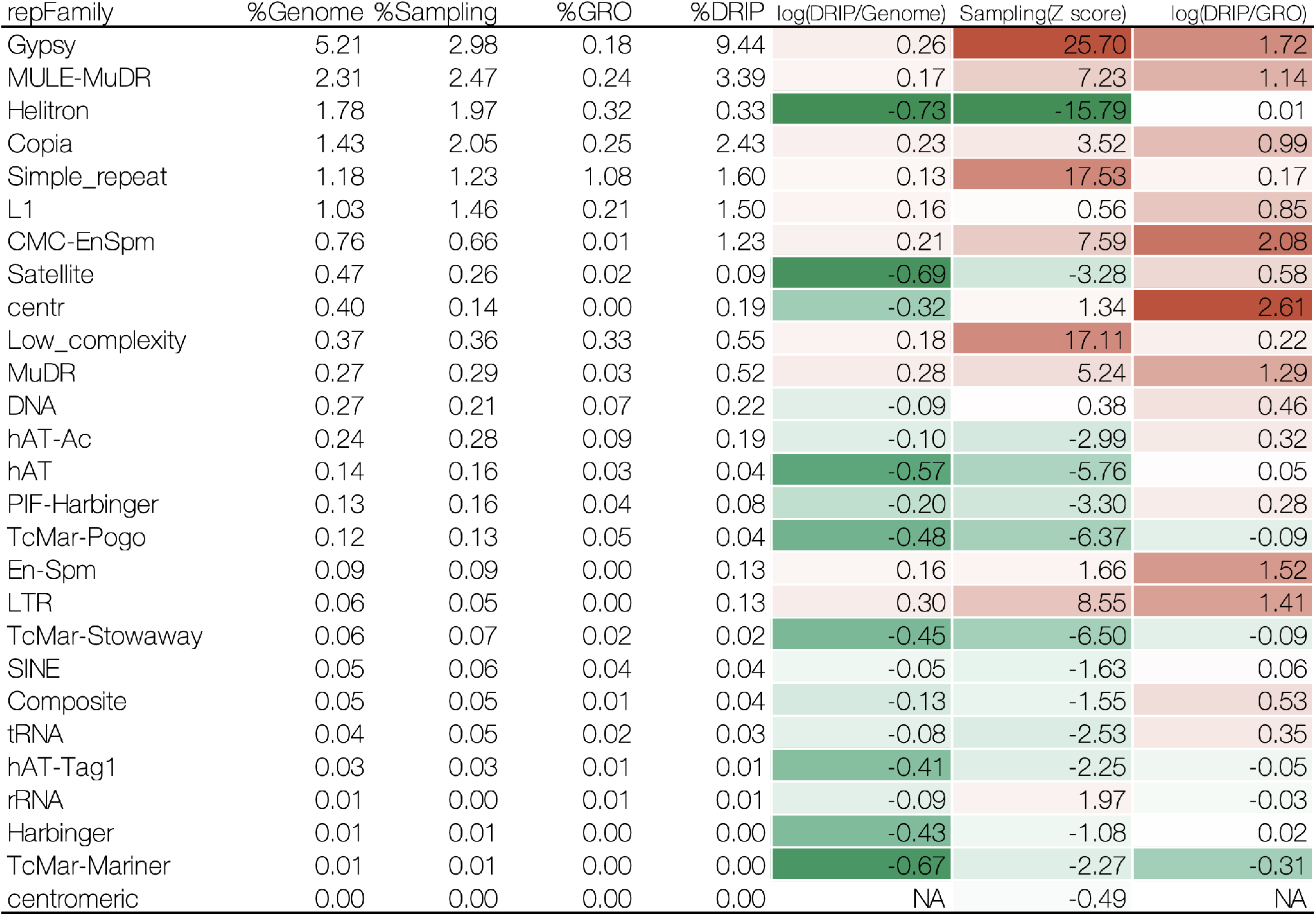
Enrichment analysis of repeat families in A. thaliana seedlings.

## Discussion

### R-loops and transcribed regions are not necessarily overlapping

We utilized GRO-seq data to define transcribed regions for comparison with R-loop-forming regions. This is because previous reports have shown that R-loop formation is associated with transcribed regions^3,24,44^. Unexpectedly, we found that the transcribed regions and R-loops do not inevitably overlap and differ among species and developmental stages (Figure 2). We speculate that there may be several reasons for this observation. First, different experimental conditions might result in inconsistency in the GRO-seq and DRIP-seq data, since we combined them from distinct studies. Second, as mentioned above, DRIP-seq signals that do not overlap with GRO-seq might contain transforming R-loops. Third, stable and long-lived RNAs that are not detectable with GRO-seq might form R-loops. The logic behind this may be that GRO-seq is more sensitive to nascent RNA, whereas those RNAs that have been transcribed to be stable in the R-loops are undetectable. Additionally, a large fraction of R-loops, especially in fly embryos, was derived from genic regions but did not overlap with the GRO-seq data. This might indicate that some of the genes are transcribed and stably form an R-loop in cis, suggesting that R-loop formation might not be necessarily associated with transcribed genes. These results imply that the variety of R-loop formations may be more diverse than previously expected. Further experiments with the same conditions are required to investigate these hypotheses.

### Repetitive elements differently contribute to R-loop formation and function

We found that repetitive elements contribute differently to R-loop formation among the samples investigated in this study. human U2OS cells showed that 21.19% of the DRIP-seq signals contained repetitive elements, while 43.19% of the GRO-seq data contained these signatures (Figure S1A). Further analysis revealed that some repetitive sequences, especially TEs, including LINEs, SINEs, LTRs, and DNA families, did not tend to form R-loops in U2OS cells. On the other hand, low-complexity, satellite, simple repeat, retroposon, snRNA, and rRNA sequences were enriched in R-loop regions compared to non-repetitive sequences in the GRO-seq control. These results are consistent with previous reports^3,24,44,46,47^. Notably, a recent report has shown that low-complexity and simple repeat sequences are strongly associated with promoter regions^48^, as are R-loop structures^3,24,44^. These results suggest that repetitive elements, such as low-complexity and simple repeats, are the key features of R-loop formation in promoter regions. Interestingly, low-complexity sequences have also been shown to be associated with Ezh2 binding, which is a component of polycomb repressive complex 2 (PRC2), and have methyltransferase activity for histone H3 lysine 27^48^. Another report has shown that R-loop formation is required for the recruitment of PRC2 and repression of a subset of polycomb target genes^11^. These results suggest that R-loop formation involving low-complexity elements could be important for the recruitment of PRC2 and epigenetic regulation of target genes. Therefore, we hypothesize that repetitive elements in R-loop regions might contribute differently to the subsequent function of R-loop formation.

In contrast to human U2OS cells, *A. thaliana* seedlings showed that 22.25% of the DRIP-seq signals contained repetitive elements, while only 3.08% of the GRO-seq data contained these elements (Figure S1C). In addition to simple repeats, low-complexity, and satellites, which are prone to form R-loops in human U2OS cells, TEs, including LTRs, DNA transposons, and LINEs, were more preferentially enriched in R-loop regions in *A. thaliana* seedlings. These results imply that R-loop formation does not simply depend on genomic sequence features but depends highly on the species (or biological contexts). Given that R-loop formation is essential for epigenetic regulation^3,24,44^, TEs that form R-loops could be critical regulatory elements for gene regulation in *A. thaliana* seedlings. Further analysis of such factors will reveal the functional significance of R-loop formation in TEs.

To investigate the contribution of repetitive elements in R-loop formation at different developmental stages, we compared the distribution of repetitive elements in R-loop regions between fly embryos and S2 cell lines. In fly embryos, 15.5% of the DRIP-seq signals contained repetitive elements, as compared to only 5.35% of the GRO-seq data (Figure S1B). In S2 cells, 9.81% of the DRIP-seq signals contained some repetitive elements, while 6.28% of the GRO-seq data contained those elements (Figure S1B). These results show that repetitive element contribution to R-loop formation is more prominent in embryos than in S2 cells, suggesting that the impact of repetitive elements on R-loop formation remarkably changes in different developmental stages or cell lineages. We also observed that LTRs, LINEs, and satellites were highly enriched in embryo R-loops and were less enriched in S2 R-loops. Conversely, simple repeats and low complexity were relatively enriched in S2 cells and less enriched in embryos. We speculate that repetitive elements could change their function through R-loop formation, along with the developmental context. For example, gypsy, which is known as one of the major insulator elements in flies^49^, is more highly enriched in embryo R-loops than S2 R-loops. R-loop formation on gypsy may alter the function of the insulator or protein complex on insulator bodies, resulting in the downstream regulation of the chromatin compartment. This case is consistent with the recent observation that R-loop formation is associated with an enhancer- and insulator-like state^3^. Further investigation is required to reveal the relationship between R-loop formation and the insulator function of gypsy elements.

### R-loop formation might be derived from TE regulation

Our results highlight the impact of TE elements on R-loop formation, especially at different developmental stages. This suggests that the TE sequence itself could tend to form an R-loop. Because TEs originate from exogenous viruses, they are the target of gene silencing by multiple layers of defense mechanisms to prevent the harmful effects of TE activity. Therefore, R-loop formation involving TEs might be one such mechanism by which cells mitigate the effects of TEs. It has been shown that R-loop formation can stimulate transcription of an antisense sequence, resulting in the formation of heterochromatin^50,51^. This mechanism is suitable if R-loop formation has a role in silencing TE elements. Similarly, it is reasonable that R-loops have a role in regulatory signals of epigenetic regulations if their functional origin is derived from TE regulation. Moreover, chromatin loosening following the depletion of histone H1 induces the accumulation of R-loops in heterochromatic regions enriched with repetitive elements, including several types of TEs^52^. This result suggests that TE elements could preferentially form R-loop structures, when their silencing by heterochromatin is resolved. This is consistent with the notion that transcribing TE sequences increase the likelihood of R-loop formation. Taken together, R-loop formation might be intimately correlated with TE sequences, although further experimental studies are required to confirm this hypothesis.

## Conclusion

In this study, we reanalyzed DRIP-seq data to investigate the impact of repetitive elements on R-loop formation. We found that satellites, LINEs, and DNA were enriched for R-loops in humans, fruit flies, and *A. thaliana*, respectively. Consistently, we observed that R-loops preferred to form in regions of low-complexity or simple repeats across species. Additionally, we also found that the repetitive elements associated with R-loop formation differ according to the developmental stage. LINEs and LTRs are more likely to promote R-loop formation in embryos (fruit fly). While R-loop formation changes in S2 cells to being more prevalent in low complexity and simple repeat areas of the genome. These results imply that repetitive elements may have species-specific or development-specific regulatory effects on R-loop formation. To our knowledge, this is the first study to analyze the association between repetitive elements and R-loop formation across species and developmental stages. Our results show that various repetitive elements may distinctly contribute to R-loop formation in a biological context-dependent manner. This work advances our understanding of repetitive elements and R-loop biology; future research should aim to determine the mechanism of R-loop formation on each repetitive element and its biological function.

## Materials and Methods

Genomic sequences and gene annotations of human (hg38) and fruit fly (dm6) were downloaded from the UCSC genome browser^53^. The *A. thaliana* genome (TAIR10) and gene annotation were obtained from Ensembl^54^. Repetitive sequences were downloaded from the UCSC repeatmasker track^53^. We extracted genome-wide R-loop regions and transcribed regions from DRIP-seq and GRO-seq (global run-on sequencing) data, respectively, following the steps mentioned below. Bedtools (v2.25.0)^55^ were used to extract (intersect sub-command) or remove (subtract sub-command) overlaps between regions. Statistics and enrichment analysis were implemented in home-made Python scripts.

### DRIP-seq analysis

Reads were aligned to the corresponding genome using BWA-MEM (0.7.17-r1188)^56^ with default parameters. For paired-end reads (fruit fly and *A. thaliana*), reads aligned in a proper pair (SAMtools view -f 2)^57^ were considered as mapped reads. For single-end reads (human), we extracted mapped reads (SAMtools view -F −4) for subsequent analysis. Mapped reads were sorted by SAMtools, and polymerase chain reaction (PCR) duplicates were marked with Picard MarkDuplicates (v2.18.1)^58^ using default parameters. Subsequently, we discarded the read duplicates from the alignments (SAMtools view -h -F 1024).

We used DRIP-seq data from IP (immunoprecipitation), Input, and RnaseH-treated samples, separately. Using the input sample as a control, we extracted peaks in the IP and RnaseH-treated samples. MACS (v2.2.7.1) was applied to detect peaks from the alignments. In addition to the same MACS parameters (-q 0.001 -broad -broad-cutoff 0.001 -keep-dup all), we used -g hs -f BAM, -g dm -f BAMPE, and -g 1.36e8 -f BAMPE for human, fruit fly, and *A. thaliana*, respectively. We selected an output file (.broadPeak) for the final peak detection results. After removing the peaks in the IP sample that overlapped with the peaks in the RnaseH-treated sample, we obtained the final R-loop peaks.

### GRO-seq analysis

For short reads with lengths of less than 100 nt (human and fruit fly), we sequentially utilized BWA-ALN and BWA-SAMSE with default parameters for mapping. Alternatively, for *A. thaliana*, we used BWA-MEM using default parameters. Homer (v4.11)^59^ (findPeaks -style groseq) was applied to detect peaks (indicating transcripts). In cases where a tissue or cell line corresponded to multiple samples (e.g., biological replicates), we used BEDtools (merge sub-command) to combine peaks from all samples.

### Enrichment analysis

To investigate the enrichment of repetitive elements in the R-loop-forming regions, we calculated the percentage of bases in the repetitive elements in the R-loops. For this purpose, we also prepared three control groups based on different hypotheses. First, assuming that R-loops are randomly distributed in the genome, we calculated the content of the repetitive elements in the genome as a control (referred to as “genome control”). Second, assuming that R-loops have a preferential distribution of length and genomic locations, we randomly selected 1000 groups of sequences from the genome while maintaining the same number of R-loops as well as length and genomic location (referred to as “sampling control”). Finally, assuming that R-loops are overwhelmingly formed directly where they are transcribed, we used the transcribed regions defined in the GRO-seq data as a control (referred to as “GRO control”).

For genome and GRO controls, we calculated the percentage of the bases of the repetitive elements in R-loops and control sequences *x_k_* and *y_k_*, respectively, and then computed the

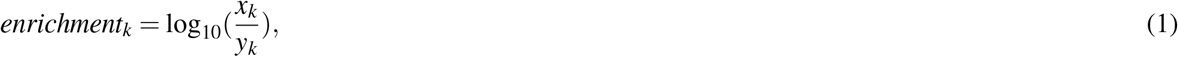

where *k* denotes a repetitive element (repeat class or family). For the sampling control, we computed the percentage of the bases of repetitive elements in R-loops and control groups as *p_k_* and {*q_k, j_*} (*j* ∈ {1*,…,* 1000}), respectively. Then, we calculated the Z score of *p_k_* in {*q_k, j_*} ( *j* ∈ {1*,…,* 1000}) as the enrichment metric.

## Availability of data and materials

DRIP-seq and GRO-seq datasets are available from public repositories. See Table S2 for the full list of the datasets.

## Acknowledgements

We thank Dr. Martin Frith and Dr. Yutaka Saito for their helpful discussions. We thank Ms. Risa Maemura for her participation in the initial survey of R-loop-related studies. Computations were partially performed on the NIG supercomputer at ROIS National Institute of Genetics.

## Author contributions

MH conceived and supervised this study. CZ designed and performed the experiments. CZ and MO wrote the manuscript. MH revised the manuscript critically. All authors contributed to analysis and interpretation of the data. All authors read and approved the final manuscript.

---

**Table S1.** Genomic distribution of DRIP- and GRO-seq peaks. Percentages of peaks overlapping with gene, promoter2k and intergenic regions shown in the last three columns. Peaks overlapping by at least 1nt with other regions are counted according to the priority (promoter2k > gene > intergenic). Promoter2k means 2000nt upstream of a gene region.

| Dataset | #Peaks | #Bases | %Gene | %Promoter2k | %Intergenic |
| --- | --- | --- | --- | --- | --- |
| U2OS(DRIP) | 10378 | 9772584 | 85.11 | 6.7 | 8.19 |
| U2OS(GRO) | 66659 | 322496237 | 86.71 | 2.86 | 10.43 |
| Embryo(DRIP) | 7354 | 7060054 | 77.37 | 5.78 | 16.86 |
| Embryo(GRO) | 7279 | 19304717 | 91.8 | 5.02 | 3.18 |
| S2(DRIP) | 12049 | 9815359 | 75.87 | 5.64 | 18.49 |
| S2(GRO) | 5812 | 42665805 | 90.79 | 4.38 | 4.83 |
| Seedling(DRIP) | 14809 | 20392020 | 63.26 | 13.57 | 23.17 |
| Seedling(GRO) | 12485 | 57831988 | 84.85 | 9.63 | 5.52 |

**Table S2.** DRIP-seq and GRO-seq datasets used in this study.

| DRIP-seq |  |  |  |
| --- | --- | --- | --- |
| Species | Source | File Name | URLs |
| Human | U2OS | Villarreal2020_U2OS_IP_1-1 | https://sra-download.ncbi.nlm.nih.gov/traces/sra64/SRR/012121/SRR12412380 |
|  |  | Villarreal2020_U2OS_IP_1-2 | https://sra-download.ncbi.nlm.nih.gov/traces/sra51/SRR/012121/SRR12412381 |
|  |  | Villarreal2020_U2OS_IP_1-3 | https://sra-download.ncbi.nlm.nih.gov/traces/sra58/SRR/012121/SRR12412382 |
|  |  | Villarreal2020_U2OS_IP_2-1 | https://sra-download.ncbi.nlm.nih.gov/traces/sra77/SRR/012121/SRR12412383 |
|  |  | Villarreal2020_U2OS_IP_2-2 | https://sra-download.ncbi.nlm.nih.gov/traces/sra57/SRR/012121/SRR12412384 |
|  |  | Villarreal2020_U2OS_Input-1 | https://sra-download.ncbi.nlm.nih.gov/traces/sra50/SRR/012121/SRR12412385 |
|  |  | Villarreal2020_U2OS_Input-2 | https://sra-download.ncbi.nlm.nih.gov/traces/sra57/SRR/012121/SRR12412386 |
|  |  | Villarreal2020_U2OS_Input-3 | https://sra-download.ncbi.nlm.nih.gov/traces/sra59/SRR/012121/SRR12412387 |
|  |  | Villarreal2020_U2OS_RH_1-1 | https://sra-download.ncbi.nlm.nih.gov/traces/sra45/SRR/012121/SRR12412388 |
|  |  | Villarreal2020_U2OS_RH_1-2 | https://sra-download.ncbi.nlm.nih.gov/traces/sra63/SRR/012121/SRR12412389 |
|  |  | Villarreal2020_U2OS_RH_2-1 | https://sra-download.ncbi.nlm.nih.gov/traces/sra71/SRR/012121/SRR12412390 |
|  |  | Villarreal2020_U2OS_RH_2-2 | https://sra-download.ncbi.nlm.nih.gov/traces/sra55/SRR/012121/SRR12412391 |
|  |  | Villarreal2020_U2OS_RH_2-3 | https://sra-download.ncbi.nlm.nih.gov/traces/sra38/SRR/012121/SRR12412392 |
| Fruit fly | Embryo | Alecki2020_Embryo_IP_1 | ftp://ftp.ddbj.nig.ac.jp/ddbj_database/dra/sralite/ByExp/litesra/SRX/SRX544/SRX5443751/SRR8645632/SRR8645632.sra |
|  |  | Alecki2020_Embryo_IP_2 | ftp://ftp.ddbj.nig.ac.jp/ddbj_database/dra/sralite/ByExp/litesra/SRX/SRX544/SRX5443752/SRR8645633/SRR8645633.sra |
|  |  | Alecki2020_Embryo_RH_1 | ftp://ftp.ddbj.nig.ac.jp/ddbj_database/dra/sralite/ByExp/litesra/SRX/SRX544/SRX5443753/SRR8645634/SRR8645634.sra |
|  |  | Alecki2020_Embryo_RH_2 | ftp://ftp.ddbj.nig.ac.jp/ddbj_database/dra/sralite/ByExp/litesra/SRX/SRX544/SRX5443754/SRR8645635/SRR8645635.sra |
|  |  | Alecki2020_Embryo_Input_1 | ftp://ftp.ddbj.nig.ac.jp/ddbj_database/dra/sralite/ByExp/litesra/SRX/SRX544/SRX5443755/SRR8645636/SRR8645636.sra |
|  |  | Alecki2020_Embryo_Input_2 | ftp://ftp.ddbj.nig.ac.jp/ddbj_database/dra/sralite/ByExp/litesra/SRX/SRX544/SRX5443756/SRR8645637/SRR8645637.sra |
|  | S2 | Alecki2020_S2_IP_1 | ftp://ftp.ddbj.nig.ac.jp/ddbj_database/dra/sralite/ByExp/litesra/SRX/SRX544/SRX5443763/SRR8645644/SRR8645644.sra |
|  |  | Alecki2020_S2_IP_2 | ftp://ftp.ddbj.nig.ac.jp/ddbj_database/dra/sralite/ByExp/litesra/SRX/SRX544/SRX5443764/SRR8645645/SRR8645645.sra |
|  |  | Alecki2020_S2_RH_1 | ftp://ftp.ddbj.nig.ac.jp/ddbj_database/dra/sralite/ByExp/litesra/SRX/SRX544/SRX5443765/SRR8645646/SRR8645646.sra |
|  |  | Alecki2020_S2_RH_2 | ftp://ftp.ddbj.nig.ac.jp/ddbj_database/dra/sralite/ByExp/litesra/SRX/SRX544/SRX5443766/SRR8645647/SRR8645647.sra |
|  |  | Alecki2020_S2_Input_1 | ftp://ftp.ddbj.nig.ac.jp/ddbj_database/dra/sralite/ByExp/litesra/SRX/SRX544/SRX5443767/SRR8645648/SRR8645648.sra |
|  |  | Alecki2020_S2_Input_2 | ftp://ftp.ddbj.nig.ac.jp/ddbj_database/dra/sralite/ByExp/litesra/SRX/SRX544/SRX5443768/SRR8645649/SRR8645649.sra |
| *A.thaliana* | seedlings | Xu2017_Seedling_IP_1 | ftp://ftp.ddbj.nig.ac.jp/ddbj_database/dra/sralite/ByExp/litesra/SRX/SRX261/SRX2617507/SRR5318040/SRR5318040.sra |
|  |  | Xu2017_Seedling_IP_2 | ftp://ftp.ddbj.nig.ac.jp/ddbj_database/dra/sralite/ByExp/litesra/SRX/SRX261/SRX2617508/SRR5318041/SRR5318041.sra |
|  |  | Xu2017_Seedling_RH_1 | ftp://ftp.ddbj.nig.ac.jp/ddbj_database/dra/sralite/ByExp/litesra/SRX/SRX261/SRX2617509/SRR5318042/SRR5318042.sra |
|  |  | Xu2017_Seedling_RH_2 | ftp://ftp.ddbj.nig.ac.jp/ddbj_database/dra/sralite/ByExp/litesra/SRX/SRX261/SRX2617510/SRR5318043/SRR5318043.sra |
|  |  | Xu2017_Seedling_Input | ftp://ftp.ddbj.nig.ac.jp/ddbj_database/dra/sralite/ByExp/litesra/SRX/SRX286/SRX2867541/SRR5626994/SRR5626994.sra |

| GRO-seq |  |  |  |
| --- | --- | --- | --- |
| Human | U2OS | GRO_U2OS | https://www.ncbi.nlm.nih.gov/geo/query/acc.cgi?acc=GSE66928 |
| Fruit fly | Embryo | GRO_Embryo | https://www.ncbi.nlm.nih.gov/geo/query/acc.cgi?acc=GSE41611 |
|  | S2 | GRO_S2_A | https://www.ncbi.nlm.nih.gov/geo/query/acc.cgi?acc=GSE68677 |
|  |  | GRO_S2_B | https://www.ncbi.nlm.nih.gov/geo/query/acc.cgi?acc=GSE23543 |
| *A.thaliana* | seedlings | GRO_seedlings | https://www.ncbi.nlm.nih.gov/geo/query/acc.cgi?acc=GSE117014 |

**Figure S1.**
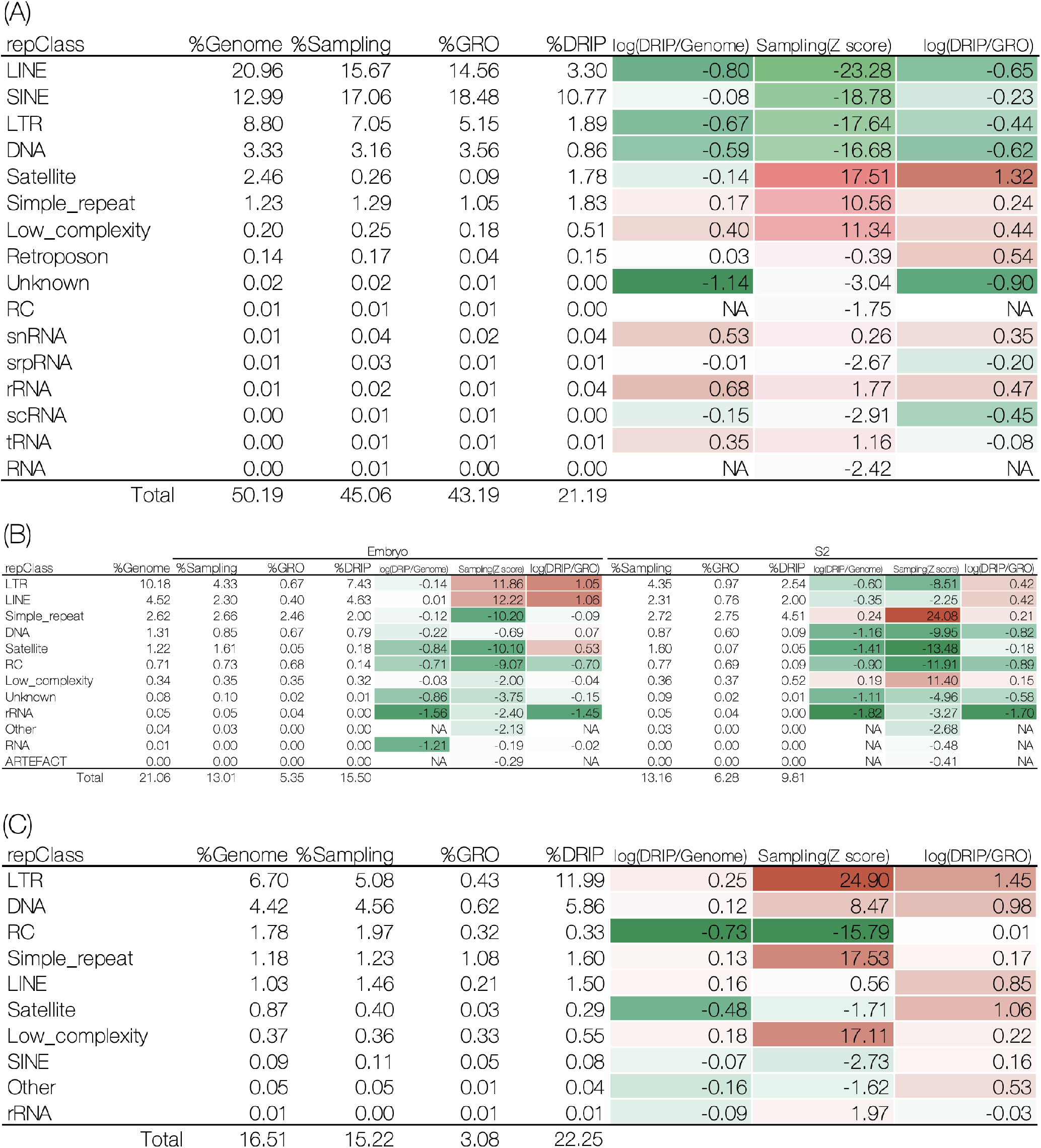
Enrichment analyses of repeat classes across (A)human, (B)fruit fly, and (C) *A. thaliana*.

